# The human cytomegalovirus protein UL116 interacts with the viral ER resident glycoprotein UL148 and promotes the incorporation of gH/gL complexes into virions

**DOI:** 10.1101/2020.11.17.387944

**Authors:** Mohammed N.A. Siddiquey, Eric P. Schultz, Qin Yu, Diego Amendola, Giacomo Vezzani, Dong Yu, Domenico Maione, Jean-Marc Lanchy, Brent J. Ryckman, Marcello Merola, Jeremy P. Kamil

## Abstract

Heterodimers of glycoproteins H (gH) and L (gL) comprise a basal element of the viral membrane fusion machinery conserved across herpesviruses. In human cytomegalovirus (HCMV), a glycoprotein encoded by *UL116* noncovalently assembles onto gH at a position similar to that occupied by gL, forming a heterodimer that is incorporated into virions. However, physiological roles for UL116 or its complex with gH remain to be identified. Here, we show that UL116 promotes the expression of gH/gL complexes and is required for the efficient production of infectious cell-free virions. *UL116-null* mutants show a 10-fold defect in production of infectious cell-free virions from infected fibroblasts and epithelial cells. This defect is accompanied by reduced expression of the two disulfide-linked gH/gL complexes that play crucial roles in viral entry: the heterotrimer of gH/gL with glycoprotein O (gO) and the pentameric complex of gH/gL with UL128, UL130, and UL131. Furthermore, gH/UL116 complexes comprise a substantial constituent of virions since an abundant gH species not covalently linked to other glycoproteins, which has long been observed in the literature, is readily detected from wild-type but not *UL116-null* virions.

Interestingly, UL116 co-immunoprecipitates with UL148, a viral ER resident glycoprotein previously shown to attenuate ER-associated degradation (ERAD) of gO, and we observe elevated levels of UL116 in *UL148-null* virions.

Collectively, our findings suggest that UL116 may serve as a chaperone for gH to support the assembly, maturation, and incorporation of gH/gL complexes into virions.

**IMPORTANCE:** HCMV is a betaherpesvirus that causes dangerous opportunistic infections in immunocompromised patients, as well as in the immune-naive fetus and preterm infants. The potential of the virus to enter new host cells is governed in large part by two alternative viral glycoprotein H (gH) / glycoprotein L (gL) complexes that play important roles in entry: gH/gL/gO and gH/gL/UL128-131. A recently identified virion gH complex, comprised of gH bound to UL116, adds a new layer of complexity to the mechanisms that contribute to HCMV infectivity. Here, we show that UL116 promotes the expression of gH/gL complexes, and that UL116 interacts with the viral ER-resident glycoprotein UL148, a factor that supports the expression of gH/gL/gO. Overall, our results suggest that UL116 is a chaperone for gH. These findings have important implications for understanding of HCMV cell tropism as well as for the development of vaccines against the virus.

## INTRODUCTION

The core viral cell entry machinery that drives membrane fusion events during herpesvirus infection is comprised of glycoprotein B (gB), and a heterodimer of glycoprotein H/ glycoprotein L (gH/gL). gB is widely posited to be the proximal viral fusogen, which undergoes a substantial conformational rearrangement during fusion of virus and target cell membranes, while gH/gL regulates the fusogenic activity of gB. Herpes simplex viruses 1 and 2 and other alphaherpesviruses express gH/gL lacking covalently bound accessory proteins. However, several different beta- and gamma-herpesviruses, such as human cytomegalovirus (HCMV), Epstein Barr virus (EBV), and human herpesviruses 6A/6B and 7, express alternative gH/gL complexes that are derivatized with virally-encoded accessory glycoproteins. At least in certain cases, the accessory glycoproteins function as receptor binding moieties. For example, the HCMV gH/gL/gO (Trimer) complex binds PDGFRαon the surface of target cells via physical interactions with gO (1–4). Meanwhile, the gH/gL/UL128/UL130/UL131 (Pentamer) complex binds neuropilin-2 (Nrp2) to drive entry into epithelial and endothelial cells, and it is the UL128-131 gene products form contacts with Nrp2 (5). The olfactory receptor OR14I1, a G-coupled protein receptor, and CD147 also are found to be required for Pentamer-dependent entry, but the mechanistic details are less clear.

Although the identification of the two alternative HCMV gH/gL complexes has been pivotal to understanding how the virus enters the broad array of cell types that the virus infects, HCMV cell tropism may also be influenced by viral gene products that modulate the relative abundance in virions of the two gH/gL complexes. For example, UL148, a viral ER resident glycoprotein, is required for high-level expression of Trimer in HCMV strain TB40/E (6), and *UL148-null* mutants of TB40/E show enhanced tropism for epithelial cells. In addition, US16 has been found to be required for incorporation of Pentamer into the virion envelope (7).

In 2016, UL116, a glycoprotein encoded by gene directly adjacent to that encoding gL *(UL115)* was reported to form a complex with gH that is found in HCMV virions (8). UL116 assembles onto gH at a position similar to that occupied by gL. However, unlike gL, UL116 does not form a disulfide linkage to gH. Furthermore, co-expression of gH with UL116 is sufficient for export of the complex from the ER, while neither glycoprotein exports efficiently on its own. UL116 therefore represents yet another component that may impact HCMV tropism, either by interacting with hitherto unidentified host cell receptors in complex with gH, or by affecting the composition of alternative gH complexes incorporated into virions.

During the course of our studies with *UL148-null* mutant HCMVs, we observed discrepancies in the ratio of gH to gL in virion preparations (see below), which led us to hypothesize that UL148 might affect the abundance of the gH/UL116 complex. Here, we present evidence that UL116 interacts with UL148, and that *UL116*-null mutants are globally defective for expression of gH/gL complexes. Our findings suggest that UL116 may act as a chaperone for gH during the assembly of gH/gL complexes.

## RESULTS

### A *UL148-null* virus lacking *UL128-131* shows mismatched ratio of gH to gL

In the course of testing whether the presence of Pentamer-specific gH/gL subunits, UL128, UL130, and UL131, might play roles in the mechanism by which UL148 regulates levels of the Trimeric gH/gL complex, gH/gL/gO, we prepared a series of mutant viruses in the background of HCMV strain TB40/E that were either (i) disrupted for *UL148* alone, (ii) ablated for the *UL128* locus, *UL128-UL131*, which encodes the three Pentamer-specific subunits, UL128, UL130, and UL131, or (iii) disrupted for both *UL148* and the *UL128* locus. Whilst analyzing the glycoprotein composition of virions produced from these mutants, we observed a puzzling discrepancy in the ratio of gH to gL in the virus lacking both the *UL128* locus (Δ128L) and *UL148* (148_STOP_)(**FIG 1**). This mutant, Δ128L_148_STOP_, showed gH levels similar to parental wildtype (WT) TB40/E, but much lower amounts of gL. The *UL148-null* virus, 148_STOP_, showed decreased levels of gH, gL, and gO, as expected (6). Whether 148_STOP_ mutant virions express a discrepant ratio of gH:gL could not be readily discerned since its levels of gH were not comparable to WT in our sample series, which were normalized for gB immunoreactivity. Because a newly identified HCMV gH complex, gH/UL116, was reported in 2016 (8), and because gH/UL116 represents a virion gH complex that lacks gL, we hypothesized that *UL148-null* virions might express higher levels of gH/UL116 compared to parental viruses that are wildtype for *UL148.* Since this novel gH complex might play roles in the hitherto poorly understood mechanisms by which UL148 influences gH composition and HCMV cell tropism, we sought to further investigate this possibility.

**FIG 1.**
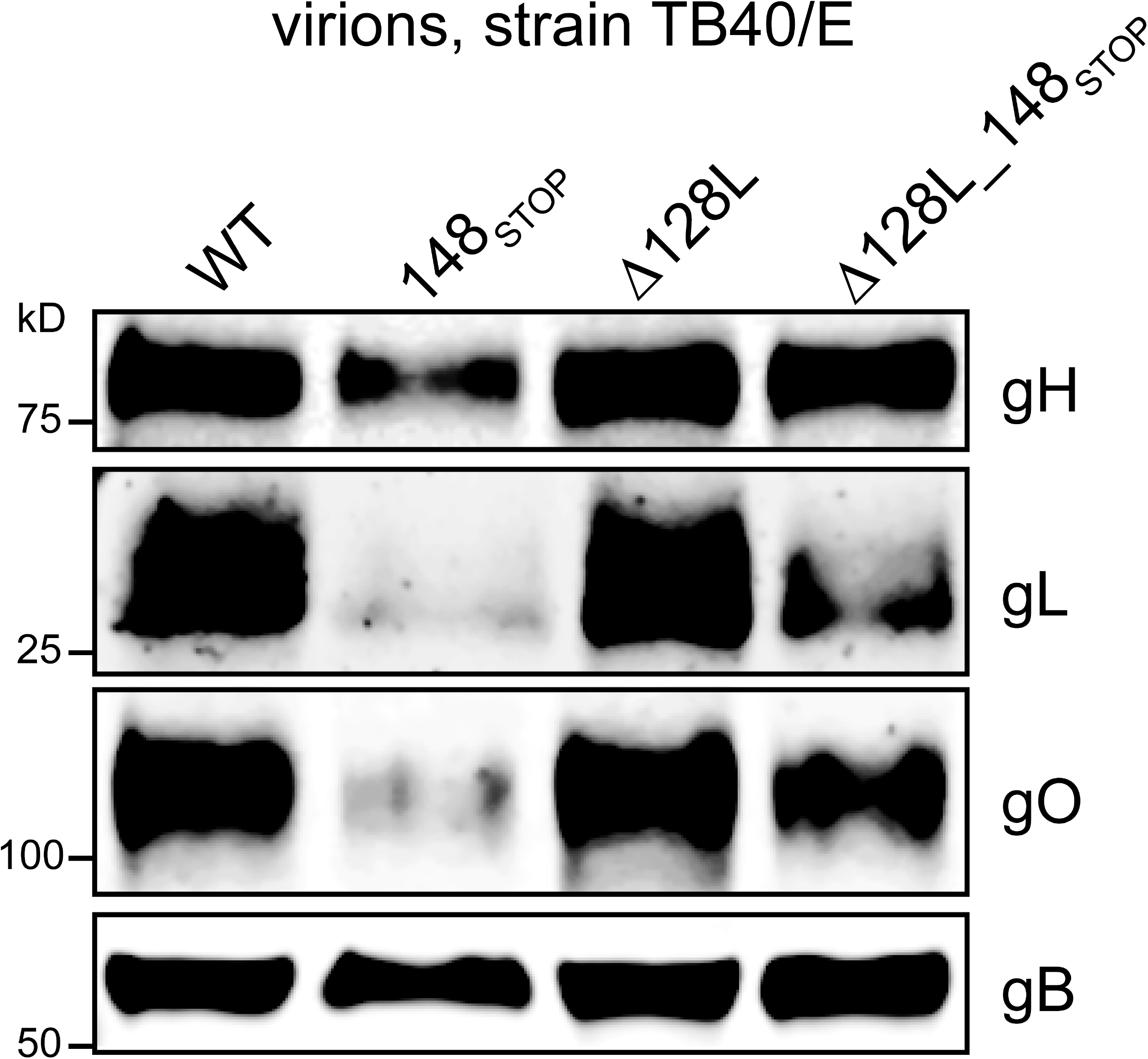
Mismatched gH:gL ratio in *UL148-null* virions. Virions produced from fibroblasts infected at MOI 1 with wildtype (WT) HCMV strain TB40/E or the indicated mutant viruses were collected at 6 days post infection, concentrated by ultracentrifugation through a sorbitol cushion, normalized for gB immunoreactivity and blotted for expression of gO, gH and gL.

### *UL116-null* viruses show a defect in production of infectious cell free virions

In order to address whether UL148 influences the abundance in virions of the gH/UL116 complex, we first constructed *UL116-null* mutant viruses since these would be important controls when investigating whether the effects of UL148 on virion gH complexes depends on UL116. We disrupted *UL116* in two different HCMV strains, AD169 and TB40/E, by replacing the 14^th^ and 15^th^ codons of the coding sequence with consecutive pair of nonsense codons. TB40/E expresses both Pentamer (gH/gL/UL128-131) and Trimer (gH/gL/gO) and retains the *UL148* allele. The AD169 derivative we used here, ADr131, carries a repaired *UL131* to rescue its expression of the Pentamer. Notably, strain AD169 does not encode a functional *UL148* and also lacks approximately 14 other viral genes due to large spontaneous deletions in the UL*Ď’* region (9).

In replication kinetics studies, the *UL116*-null viruses showed a ~1-log defect in production of cell-free infectious progeny virions following infection of human fibroblasts at MOI 1 TCID_50_ per cell (**FIG 2**). During infection of ARPE-19 epithelial cells, (again, at MOI 1, using TCID50 values as determined on fibroblasts), a similar ~1-log defect in production of infectious cell-free virions was observed for the *UL116*-null AD_r131, while the defect for *UL116*-null TB40/E was even more severe, at roughly ~2-logs. Because AD_r131 replicates more robustly in ARPE-19 than does TB40/E (6, 10), the more pronounced replication defect of *UL116*-null TB40/E compared to *UL116*-null AD_r131 in this cell type was not surprising, as we interpret this difference to most likely reflect additional rounds of replication, as might be expected when strain TB40/E viruses are used to infect ARPE-19 at MOI 1 according to titers determined in standard TCID_50_ assays, which measure infectivity in fibroblasts.

**FIG 2.**
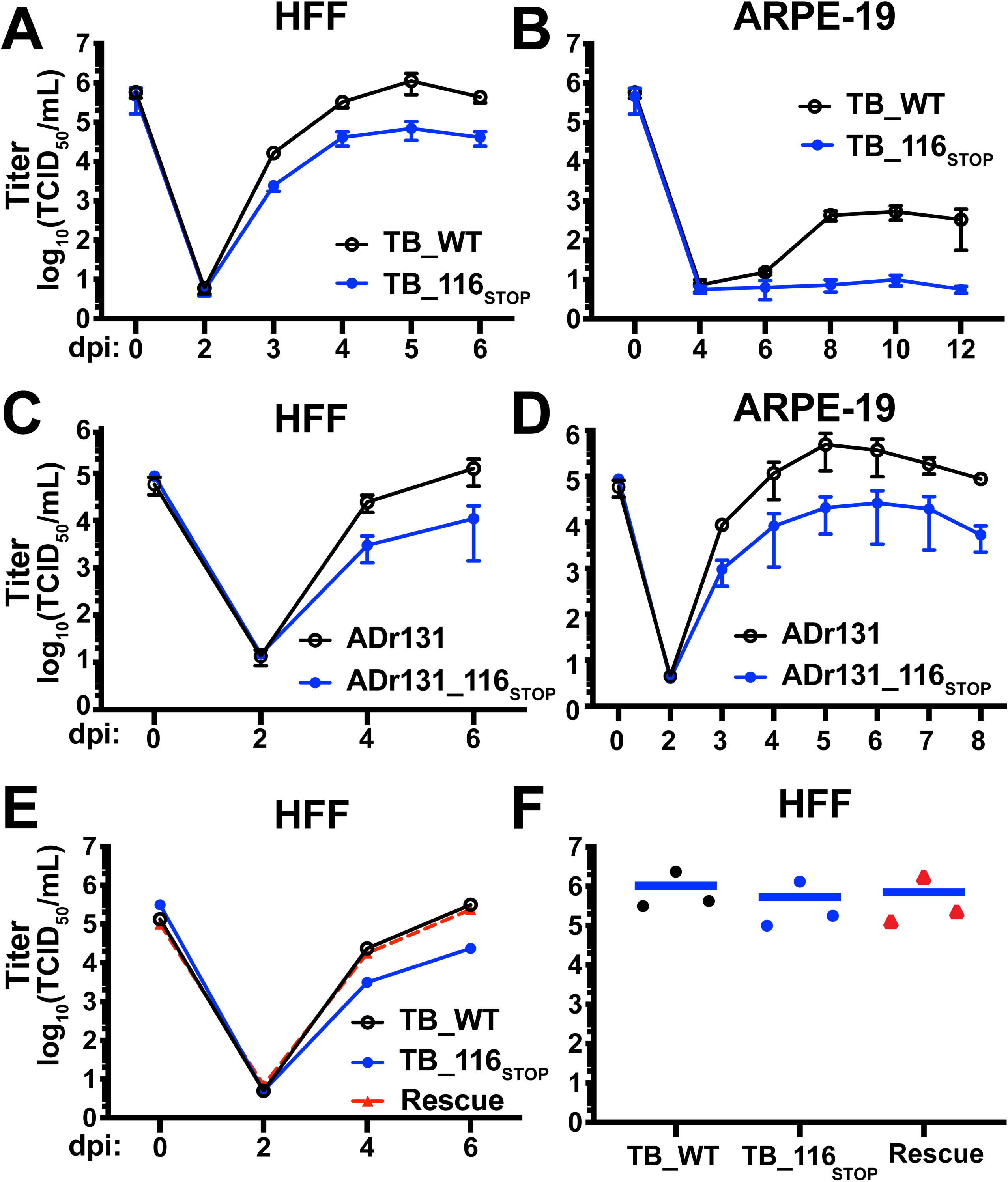
*UL116*-null mutant viruses exhibit a replication defect. (A) Fibroblasts (HFF) or (B) epithelial cells (ARPE-19) were infected at MOI 1 with strain TB40/E (TB_WT) or a *UL116*-null derivative (TB_116_STOP_). Supernatant titers from the indicated time points determined by TCID50. (C-D) Results of a similar experiment comparing wildtype vs. *UL116*-null mutant strain AD169 repaired for *UL131* (ADr131 vs ADr131_116_STOP_). (E) HFF’s were infected at MOI 1 with strain TB40/E (TB_WT), a *UL116*-null derivative (TB_116_STOP_) or a rescue of *UL116* (Rescue). (F) Cell associated virus from fibroblasts infected at MOI 1 was collected at 5 dpi and infectivity was quantified by TCID50 assay.

Importantly, a “rescuant” virus in which a functional *UL116* was restored to the *UL116*-null TB40/E mutant, replicated indistinguishably from WT parental TB40/E during infection of fibroblasts (**FIG 2E**). This finding argues against the possibility that spurious mutations elsewhere in viral genome account for the defects observed for the *UL116*-null virus. Despite its defect in production of cell-free virions, the *UL116*-null TB40/E mutant produced cell-associated titers that were similar to wildtype (**FIG 2F**). The latter finding may explain why previous genome profiling studies failed to observe a replication defect for *UL116*-null mutant viruses (11, 12). We interpret these results to suggest that UL116 is required for efficient production of infectious cell-free virions.

### *UL116-null* viruses show reduced expression of gH/gL complexes

Since UL116 is a constituent of a gH complex found in HCMV virions, we next evaluated the relative expression of the two gH/gL complexes and of gH/UL116 in infected cell lysates and in virions, as we surmised that differences in the expression of gH complexes might contribute to the differences in viral replication that we observed. After collecting virions by ultracentifugation through a sorbitol cushion, we lysed virions in non-ionic detergent to extract envelope glycoproteins, normalized gel loading such that gB levels would be equivalent across samples and carried out Western blot analyses. As expected, bands immunoreactive to an anti-UL116 monoclonal antibody were absent from *UL116*-null infected cells and from *UL116*-null virions (**FIG 3**). Interestingly, *UL116*-null mutants also showed reduced expression of gH and of other representative viral glycoproteins that participate in multiprotein complexes with gH, i.e.: gO, gL, and UL130 (**FIG 3**). Evidence of reduced expression of gH and of components of gH/gL complexes was observed in infected cell lysates and in purified virions for *UL116*-null mutants of both AD_r131 and TB40/E (**FIG 3**).

**FIG 3.**
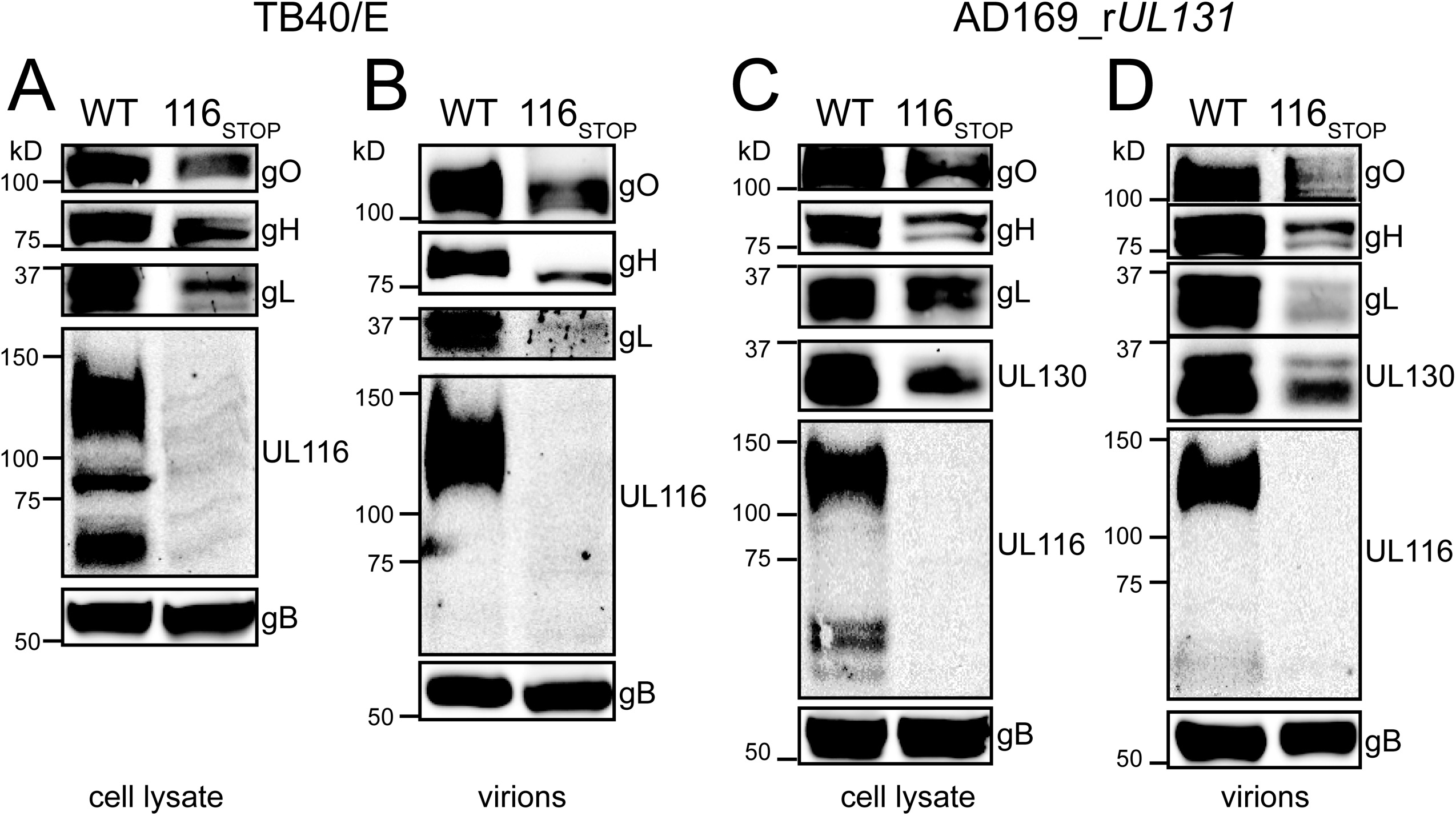
*UL116*-null viruses poorly express gH/gL complexes. (A) Lysates of HFF infected at MOI 1 with the indicated strain TB40/E derived viruses were collected at 4 dpi, resolved by SDS-PAGE, transferred to nitrocellulose and immunoblotted for gO, gH, gL, UL116, and gB. (B) Virions were collected at 6 dpi and concentrated by ultracentrifugation. Levels of gB were normalized across samples and then expression of gO, gH, gL, and UL116 were compared by Western blot. (C) Lysates of ARPE-19 cells infected at MOI 1 with the indicated strain AD169_r*UL131* derived viruses were collected at 4 dpi and immunoblotted for gO, gH, gL, UL130, UL116, and gB. (D) Virions were collected at 6 dpi and concentrated by ultracentrifugation. Levels of gB were normalized across samples and then expression of gO, gH, gL, UL130, and UL116 were compared by Western blot.

### An inhibitor of ERAD normalizes levels of gL in HCMV mutants disrupted for UL116

Although there are no known upstream splice junctions on the gL coding strand that might trivially explain a global reduction in gH/gL complexes in our *UL116*-null mutant viruses [(13), (Lars Dölken, personal communication)], since we engineered nonsense codons early in the *UL116* ORF, we wished to assure ourselves that the reduced expression of gL polypeptide did not reflect a failure to transcribe gL mRNA, which is encoded by *UL115*, a gene neighboring *UL116.*Because this region of the viral genome is transcriptionally complex and a number of neighboring genes share a co-terminal polyadenylation signal with *gL* mRNA, we opted to instead ask whether gL was post-translationally unstable during infection in the absence of UL116. We therefore turned to kifunenesine (KIF), a potent and specific inhibitor of alpha-mannosidase activities that play roles in marking terminally misfolded glycoproteins for ER-associated degradation (ERAD)(14). We reasoned that if loss of UL116 expression post-translationally destabilizes the assembly of gH/gL complexes, then gL levels should be rescued by KIF treatment, whereas if the *gL (UL115)* transcript were somehow being subjected to nonsense mediated decay or otherwise poorly transcribed due to the incorporation of nonsense codons in *UL116*, KIF would fail to show any effect.

KIF treatment dramatically restored gL levels in *UL116*-null mutant viruses, both in strain TB40/E and in AD169 (**FIG 4**), while failing to affect levels of gB or IE1. Reassuringly, KIF treatment did slightly increase the mobility of gB and gL, consistent with its known effects in preventing conversion of high-mannose N-glycans to more complex, higher molecular weight species. Overall, these results indicated to us that gL is likely being translated at equivalent levels in the absence of UL116, but that the protein is being targeted for ERAD to a greater degree in the *UL116*-null setting relative to WT. From these results, we conclude that differences in posttranslational stability of gL, and not polar effects of inserting nonsense codons in the *UL116* ORF, likely account for the defects that we observed in gL expression.

**FIG 4.**
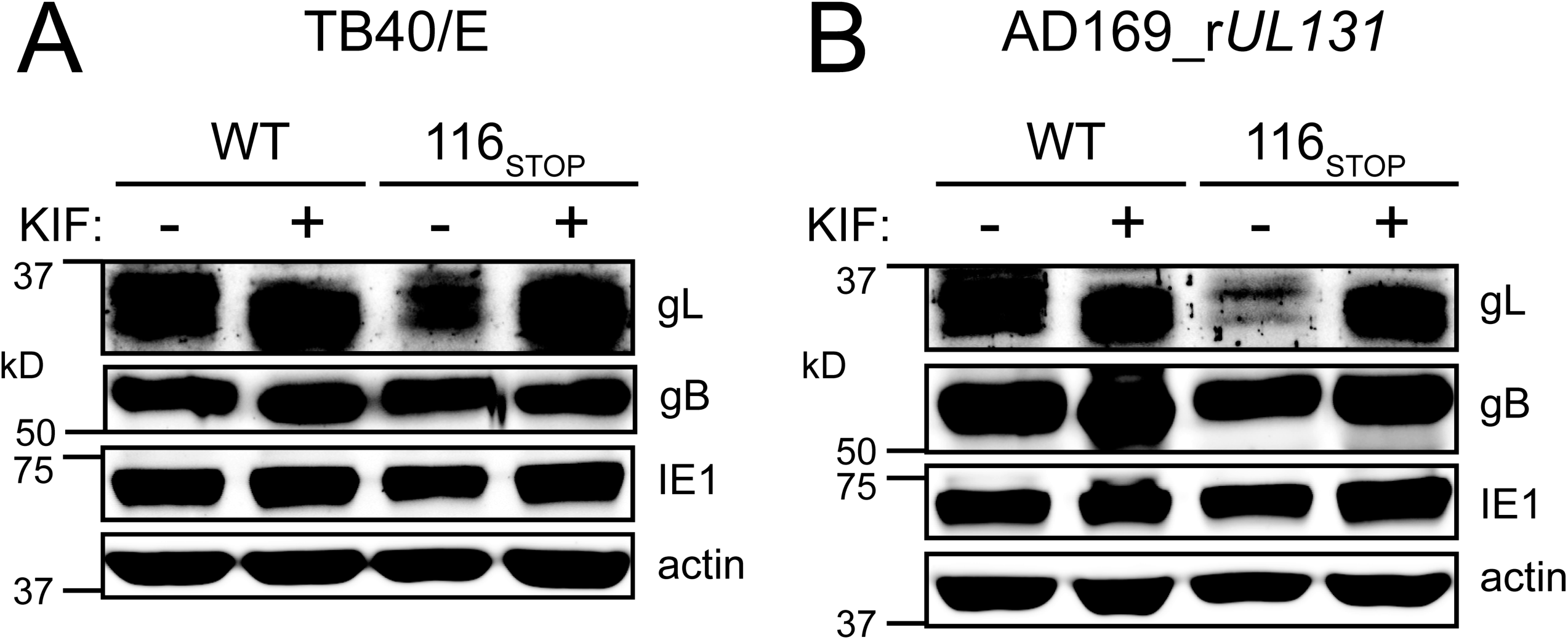
Inhibition of ERAD stabilizes gL in *UL116-null* mutant HCMVs. Fibroblasts (HFFT) were infected at MOI 1 with strain (A) TB40/E (WT) or a *UL116*-null derivative (116_STOP_), (B) AD169_r*UL131* (WT) or a *UL116*-null derivative (116_STOP_). Cells were treated with kifunensine (KIF) at 2.5 μM [final] or 0.1% carrier alone (water) at 48 hpi. At 72 hpi, expression of gL, gB, IE1 and actin levels were analyzed by Western blotting.

To assess if the observed effects of UL116 on gH/gL expression were dependent on other viral factors, we used an adenovirus (Ad) vectors to express combinations of these proteins in both fibroblasts and epithelial cells. For these experiments, we expressed a soluble gH (sgH), lacking its C-terminal tranmembrane anchor and cytoplasmic tail, from an Ad vector similar to one we used in previous studies (15). As expected (15), sgH was not secreted when expressed alone, but co-expression with gL resulted in accumulation of both proteins in the culture supernatants as apparent glycoforms of sgH/gL disulfide-linked heterodimers and homodimers of sgH/gL heterodimers (**FIG 5**), as observed in previous studies (16, 17). When UL116 was expressed in place of gL, sgH was detected in the culture supernatants as apparent sgH monomers. Co-expression of all three glycoproteins led to a drastic increase in the total amount of sgH in the supernatants, which predominately accumulated as homodimers of sgH/gL heterodimers. These results are consistent with the notion that UL116 protein acts to stabilize gH in the ER, and thereby causing increased expression of gH/gL heterodimers, which like gH/UL116 but not gH alone, are competent to can traffic toward the Golgi and beyond.

**FIG 5.**
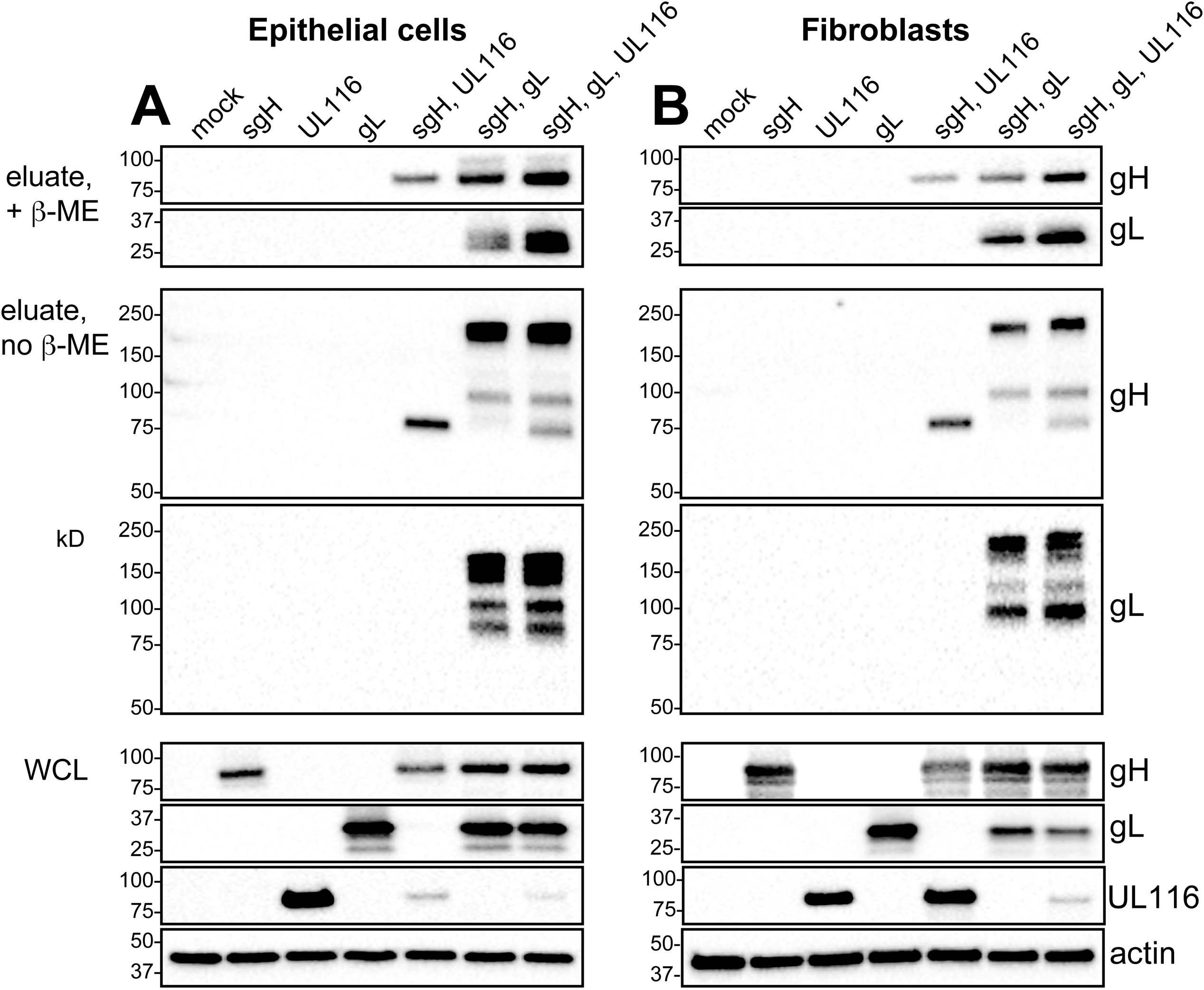
UL116 promotes enhances expression of gH/gL during ectopic coexpression of the proteins. (**A**) ARPE-19 epithelial cells or (**B**) human fibroblasts were infected with the indicated combinations of adenovirus (Ad) vectors expressing gL, UL116, or a soluble gH (sgH) fused to a C-terminal polyhistidine tag. An equal total adenovirus MOI was maintained across conditions by including a control adenovirus vector expressing eGFP. Soluble gH complexes were collected from culture supernatants using Ni-NTA agarose beads, resolved by reducing and non-reducing SDS-PAGE, and analyzed by Western blot using anti-gH or gL antibodies, as indicated.

### *UL148-null* virions incorporate increased amounts of UL116

The global decrease in the abundance of all gH-containing complexes that we observed for *UL116*-null mutants was reminiscent of *UL148*-null mutants, which likewise show decreased expression of gO and hence, deceased levels of the gH/gL/gO Trimer in virions (6). Therefore, we directly compared the expression of gH/gL complexes in isogenic null mutants of *UL116* and *UL148* in the context of strain TB40/E. As expected, *UL148*-null virus infected fibroblasts showed decreased expression of gO and gL relative to WT infected cells. *UL116*-null infections likewise showed a decrease in expression of gO and gL, which was similar in magnitude to that observed for *UL148*-null infections. Interestingly, the gO species detected from the *UL116*-null sample exhibited slower relative mobility relative to the *UL148*-null setting (**FIG 6A**), perhaps due to effects of UL148 on the secretory pathway and its activation of the integrated stress response (18).

**FIG 6.**
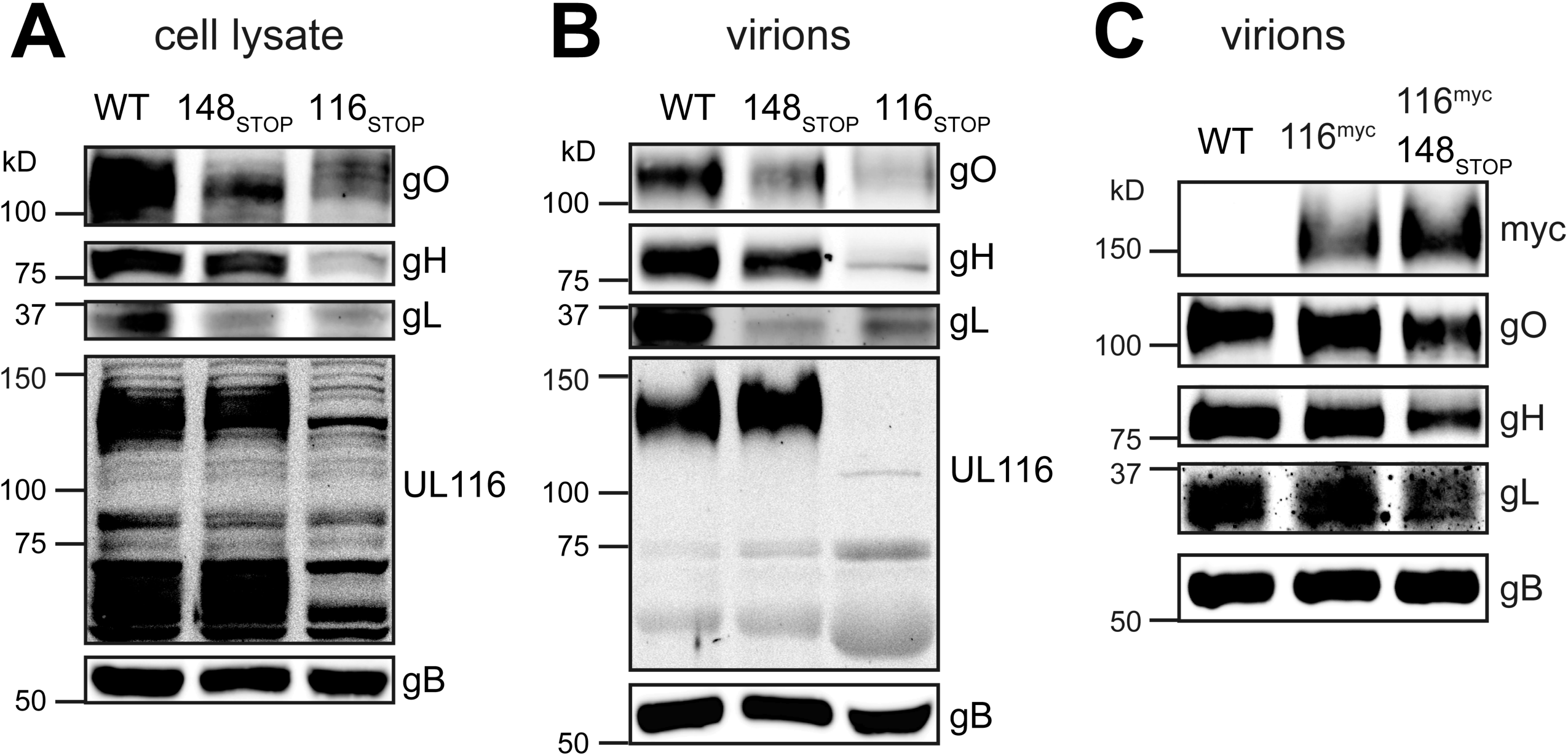
Comparison of gH/gL complexes between *UL116*-null and *UL148-null* mutants. (A) Cell lysates of fibroblasts infected at MOI 1 with the indicated HCMV strain TB40/E derived viruses were collected at 4 dpi and expression of gO, gH, gL, UL116, and gB was monitored by Western blot. (B) Virions were isolated from infected cell supernatants (collected at 6 dpi) by ultracentrifugation. After normalizing for gB loading, the levels of gO, gH, gL, and UL116 in virions were compared by Western blot. (C) 6 dpi virions were isolated by ultracentrifugation from infected cell supernatants of recombinant strain TB40/E carrying a myc tag at the C-terminus of UL116. After normalizing for gB loading, the levels of myc (UL116), gO, gH, and gL in virions were compared by Western blot.

A more obvious difference, however, is that the *UL116*-null mutant exhibited a defect in gH expression, which was not apparent in *UL148*-null setting. As would be expected, bands specifically immunoreactive to anti-UL116 monoclonal antibody (mAb) at ~130 kD and ~60 kD were absent from lysates of *UL116*-null mutant virus infected cells, while the mature virion associated ~130 kD UL116 glycoform (8) was absent from *UL116*-null virions (**FIG 6A-B**). In side-by-side comparisons of virions loaded for equivalent gB levels, gH levels were only slightly decreased in *UL148-null* virions relative to WT, while gO and gL showed a more striking decrease (**FIG 6B**). Meanwhile, the ~130 kD UL116 species appeared to be present at slightly increased levels in *UL148-null* virions relative to the WT comparator.

Results of similar experiments making use of recombinant TB40/E viruses that encode a myc epitope tag at the C-terminus of UL116 likewise indicated that *UL148-null* mutant virions show a modest, yet appreciable increase in the incorporation of UL116 (**FIG 6C**). *UL116*-null mutant virions showed poor expression of all three components of the Trimer, gO, gH, and gL, with an obvious decrease in gH abundance relative to the *UL148-null* and WT comparators (**FIG 6B**). These data suggest that the gH/UL116 complex is ordinarily a major source of gH in HCMV virions.

From these experiments, we conclude that although *UL116*-null and *UL148-null* viruses each show decreased expression of gH/gL complexes, the *UL148-null* phenotype mainly affects the abundance in virions of gO and gL, consistent with destabilized expression of the gH/gL/gO Trimer. Meanwhile, *UL148-null* mutants showed a modest-to-moderate increase in virion incorporation of gH/UL116, which would explain the compensatory effect on gH levels. However, disruption of *UL116* stabilizes the expression of all gH/gL complexes, and of course leads to an absolute loss of the gH/UL116 complex. In other words, because *UL148-null* virions carry increased levels of the gH/UL116 complex, which lacks gL, the virion levels of gL are affected more drastically than gH.

### gH/UL116 accounts for the presence in virions of a major gH species not covalently bound to other glycoproteins

Our results thus far suggest that UL116 strongly influences the abundance of gH/gL complexes in virions. Because *UL148*-null viruses produce virions that contain larger amounts of gH than would be expected given their poor incorporation of gL and gO, we hypothesized that the gH/UL116 complex is the source of the abundant virion gH species that is not covalently bound to other glycoproteins, which has long been observed in the literature (19, 20). To test this, we compared virions produced by parental WT TB40/E to those from the *UL116*-null derivative, TB_116_STOP_, and also compared virions produced by AD_r131 to its *UL116*-null derivative, AD_r131_116_STOP_, resolving virion lysates by non-reducing SDS-PAGE and then immunoblotting for gH and gL (**FIG 7**).

**FIG 7.**
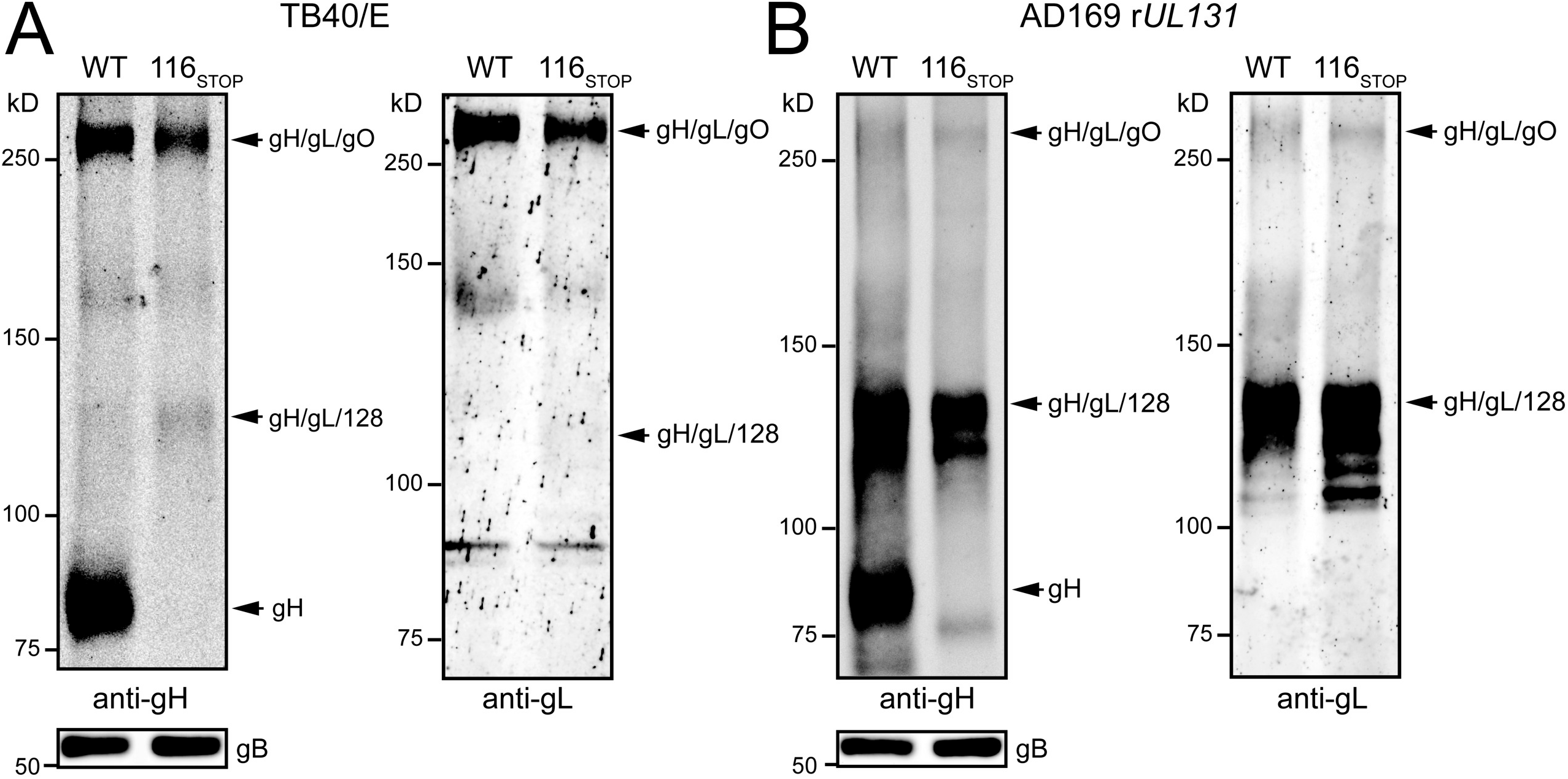
gH/UL116 accounts for the presence of an abundant virion gH species not covalently bound to other glycoproteins. Virions produced from fibroblasts infected with the indicated strain TB40/E-derived viruses were resolved by SDS-PAGE under non-reducing conditions and transferred to nitrocellulose membranes. (A) anti-gH mAb (AP-86) or (B) anti-gL polyclonal sera were used for detection in Western blots. Additionally, gB was detected by reducing SDS-PAGE to confirm comparable loading of virion material (shown below non-reducing anti-gH panels).

In virions of both strains, a prominent anti-gH immunoreactive band migrating with a relative mobility (M_r_) of ~85 kD (slightly above 75 kD molecular weight marker) was readily detected from each parental WT virus but not from the *UL116*-null mutants. As this species was not detected by anti-gL serum, we interpret it to be the gH species not covalently bound to other glycoproteins long noted to be present in HCMV virions (19, 20). From extracts of WT TB40/E virions, we readily detected the gH/gL/gO Trimer migrating at just above the 250 kD M_r_ marker, and the gH band at M_r_ ~ 85 kD, which we interpret to be monomeric gH. However, in the *UL116*-null mutant virus, we detected lower levels of Trimer, a faint band at ~135 kD matching the expected size of gH/gL/UL128, were entirely unable to detect any species matching the expected size of monomeric gH, suggesting that disruption of UL116 prevents the incorporation of gH species that are not covalently linked to other glycoproteins.

In AD_r131 virions, we observed all three of the major gH species reported by Wang and Shenk when they analyzed a similar AD169 derivative repaired for *UL131* (20) (**FIG 7B**), although the detection of the largest of these immunoreactive species, migrating just above 250 kD, which corresponds to gH/gL/gO (Trimer) was detected only faintly by both anti-gH and anti-gL sera. The latter observation is consistent with observations from studies of strains AD169 and Merlin, such as AD_r131 (20) and Merlin repaired for *UL128* (21), which suggest that the Trimer is inefficiently incorporated when Pentamer expression is high.

The immunoreactive band at M_r_ ~135 kD, interpreted to be gH/gL/UL128, was much more abundant in AD_r131 and its *UL116*-null mutant derivative than it was in TB40/E. This is consistent with findings from other groups, which suggest that TB40/E poorly expresses the Pentamer (1, 21). [The gH/gL/UL128 species is expected to lack UL130 and UL131 since these are not disulfide linked to gH/gL (20, 22, 23).] The third band detected by gH antibodies, which migrated at approximately 85 kD, was readily detected from parental AD_r131 but not *UL116*-null virus. Again, we interpreted this band to be the gH species described in the two earlier studies to be comprised of gH monomers that are not disulfide linked to other glycoproteins (19, 20).

In the *UL116*-null derivative (116_STOP_), only the largest immunoreactive band, interpreted as the gH/gL/gO Trimer, and the immunoreactive band at ~135 kD interpreted as gH/gL/UL128 were readily observed, with the Trimer band being only weakly detected, while the Mr ~85 kD band, interpreted to be gH lacking any disulfide-linked accessory proteins, was not detected. However, a faint anti-gH immunoreactive band at M_r_ 75 kD did appear in the 116_STOP_ condition (**FIG 7B**, left panel). Since this band migrates more rapidly than authentic gH, we interpreted it to be non-specific signal, although we cannot exclude the possibility that small amounts of hypoglycosylated gH were present in these *UL116*-null virion preparations.

Results from a duplicate set of samples probed with antiserum specific for gL further suggest that (i) the ~85 kD species is a gH band that lacks gL, (ii) the ~135 kD band is gH/gL/UL128, and (iii) the ~250 kD band is gH/gL/gO (**FIG 7**). The only species the gL antisera robustly detected in virions of both AD_r131 and its *UL116*-null derivative was the ~135 kD band that we interpret as gH/gL/UL128. Notably, the gL antibody also detected gH/gL/gO complex from AD_r131 virions while failing to produce signal for the ~85 kD species robustly detected by gH antibody (**FIG 7A, 7B**), which further supports our conclusion that the ~85 kD band is monomeric gH.

The species interpreted as gH/gL/UL128 appeared to be expressed in at least three glycoforms in *UL116*-null mutant AD_r131 virions, with only the slowest migrating and presumably largest glycoform being present in virions from the parental AD_r131 virus. This may indicate that the pentamer is unable to mature efficiently without UL116. Overall, from these results we conclude that *UL116*-null viruses produce virions that lack a prominent gH species found in WT virions. Further, the simplest interpretation of our data is that this species is gH/UL116.

### Virions produced from *UL116-null* infections are poorly infectious

We next sought to determine whether *UL116*-null virions of would show differences in virion infectivity relative to WT. For these studies, we compared WT strain TB40/E (TB_WT) to *UL116*-null (TB_116_STOP_) and *UL148-null*(TB_148_STOP_) derivatives. We chose to use this set of strain TB40/E viruses since TB_148_STOP_ would provide control in which tropism differences would be expected, and we did not yet know how TB_116_STOP_ would behave. Preparations of cell-free virus from infected cell supernatants were measured by qPCR to determine the number of DNAse I-resistant (presumably enveloped) HCMV genomes. Then equivalent numbers of viral genomes were measured for infectivity by TCID_50_ assay on fibroblasts and ARPE-19 epithelial cells. We found that *UL116*-null TB40/E virions were poorly infectious for both fibroblasts and epithelial cells. In contrast, TB_148_STOP_ virions showed ~2-fold reduced infectivity for fibroblasts but approximately 10-fold improved infectivity for ARPE-19 (**FIG 7**), roughly in-line with our previous findings (6). From these data, we conclude that virions released to the medium from *UL116*-null TB40/E infected fibroblasts are poorly infectious on both fibroblasts and epithelial cells.

### UL148 co-immunoprecipitates with UL116

Because UL148 and UL116 each appear to positively influence virion incorporation of one or more constituents of HCMV gH/gL complexes, we sought to determine whether the two proteins might interact. Therefore, we generated a wildtype and *UL148-null* strain TB40/E viruses that express a myc-tag at the C-terminus of UL116. We then carried out anti-myc immunoprecipitation to ask whether UL148 would co-immunoprecipitate from infected cells with UL116. Indeed, we observed a band strongly immunoreactive to UL148 antibodies was robustly detected from anti-myc immunoprecipitates of cells infected with TB_116^myc^, but not TB_148_STOP__116^myc^ (**FIG 8**). Notably, gL was not detected from anti-myc IPs of either virus infection setting, while gH was readily detected in both settings, as would be expected since UL116 has been shown to form gL- free complex with gH (8).

**FIG 8.**
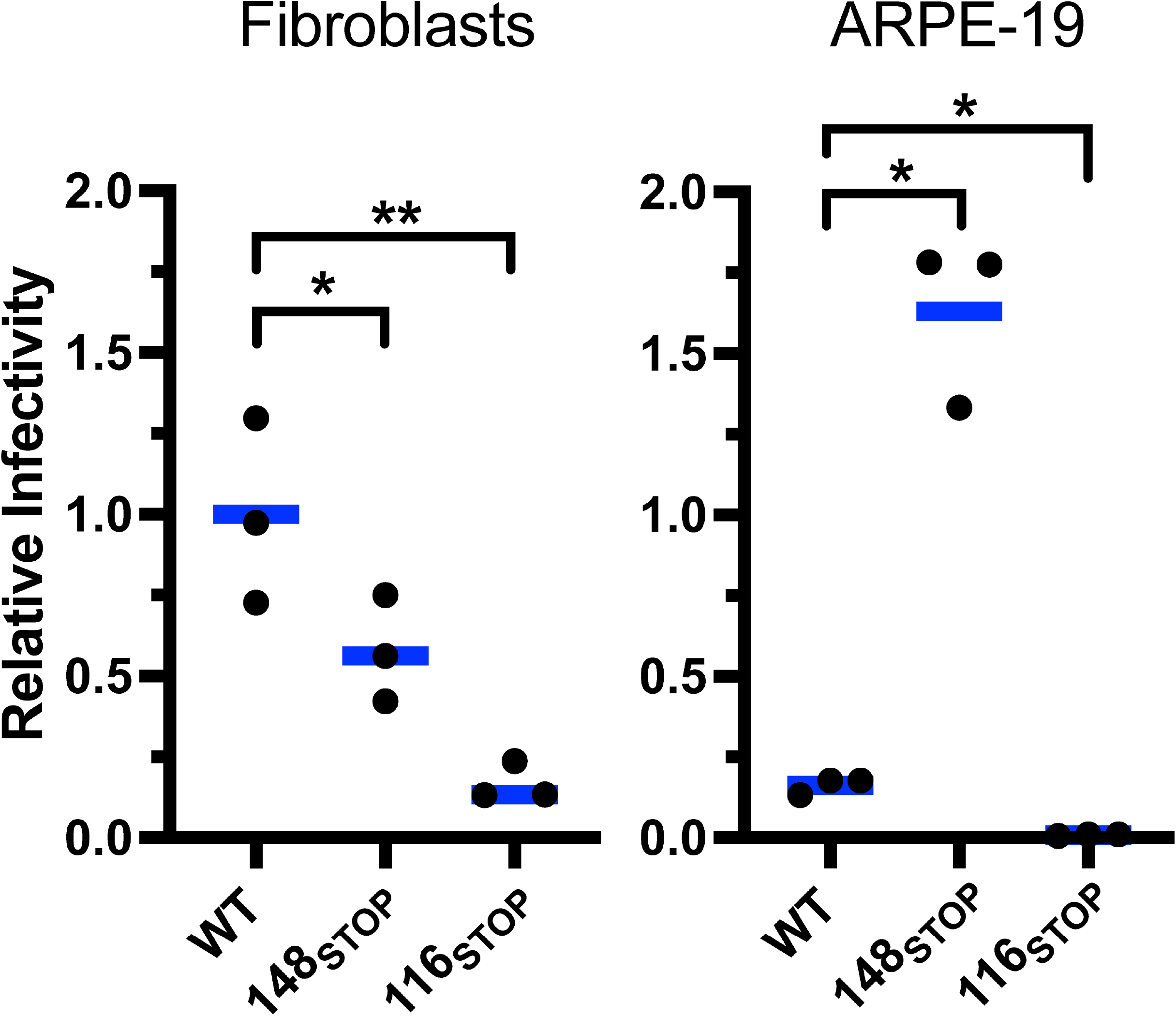
Tropism assay. Fibroblasts were infected at MOI 1 with strain TB40/E (TB_WT), a *UL148*-null derivative (TB_148_STOP_) or a *UL116*-null derivative (TB_116_STOP_). Cell free virions were quantified by qPCR. HFFs or ARPE-19 cells were infected in wells of 96-well plates at 50 genomes per cell. Cells were fixed and stained at 30 hpi using an IE1 monoclonal antibody to identify wells containing infected cells for determination of TCID50/viral genome. Relative infectivity was calculated by setting wild-type (TB_WT) infectivity on fibroblasts as 1.0 and expressing results for other conditions relative to that.

Since we did not yet have a virus carrying HA-tagged UL148 and myc tagged UL116, and because our rabbit anti-HA is not suitable for IP (not shown), we also asked whether the putative protein-protein interaction between UL116 and UL148 could be detected from the setting of cells transiently expressing the viral glycoproteins. We therefore transfected HEK-293T cells with plasmid expression vectors for gH, myc-tagged UL116 (UL116^myc^), and HA-tagged UL148 (UL148^HA^), individually and in each combination of plasmids. 48 h later, we lysed cells and immunoprecipitated using anti-HA and anti-gH antibodies.

We found that gH and UL116 detectably immunoprecipitated with HA antibodies, when constructs encoding either or both proteins were co-transfected with a plasmid expressing HA-tagged UL148 (**FIG 9**). Meanwhile, HA-tagged UL148 was readily detected in anti-gH immunoprecipitates (IPs) of cells cotransfected with expression vectors for gH and UL148^HA^, or gH, UL148^HA^ and UL116^myc^, but not when UL148^HA^ was expressed on its own, which suggested that UL148 did not promiscuously or non-specifically contaminate the anti-gH IPs. As expected, anti-gH antibodies robustly immunoprecipitated both gH and UL116^myc^ when both proteins were present. Overall, these coimmunoprecipitation (co-IP) results from infected and transfected cell contexts together suggested to us that UL148 and UL116 are either able to physically interact or participate together in a complex with other cellular and or viral glycoproteins.

**FIG 9.**
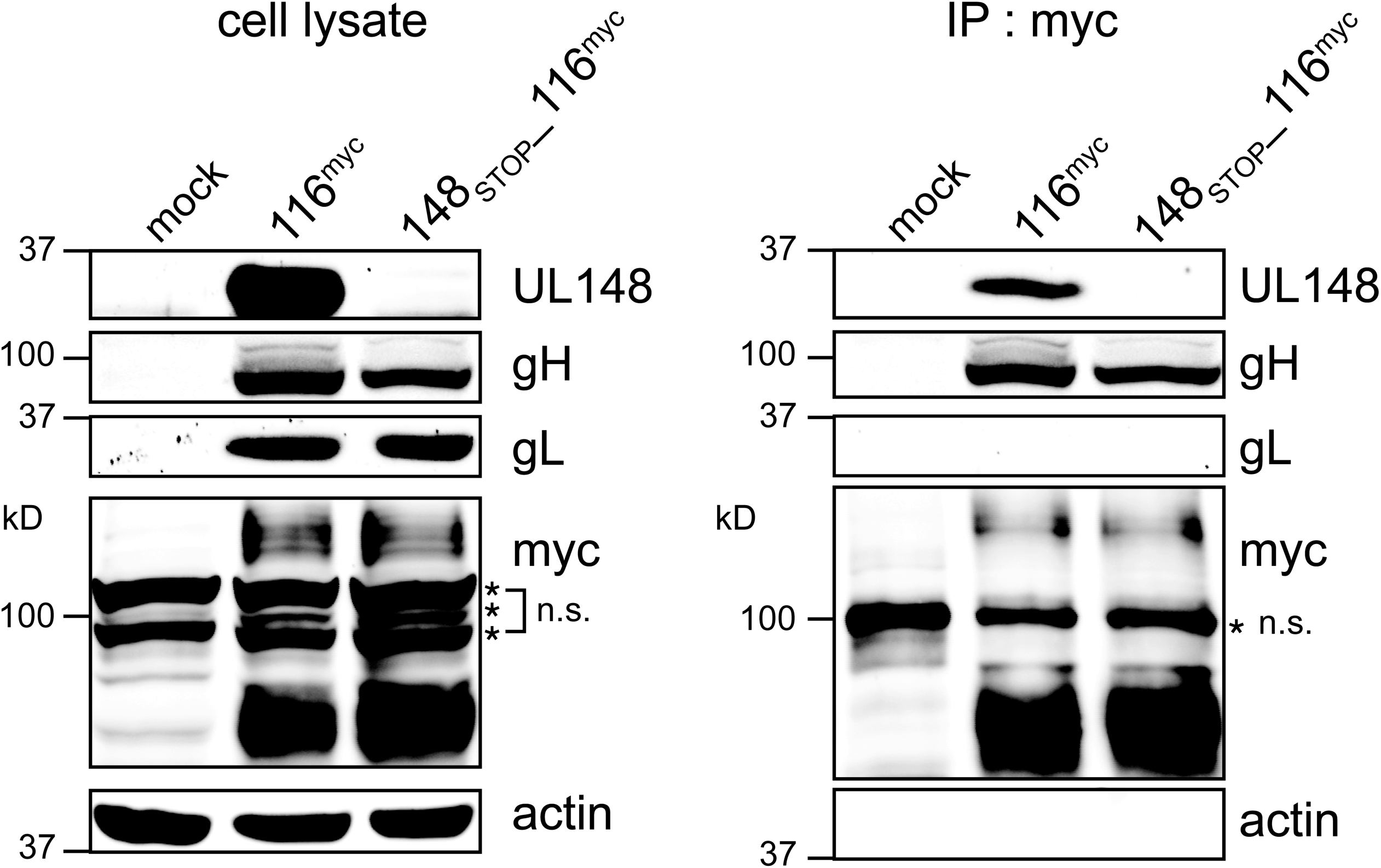
UL148 co-IPs from infected cells with UL116. Fibroblasts were infected at MOI 1 with HCMV strain TB40/E carrying a myc tag at the C-terminus of UL116 (116^myc^) or a *UL148-null* mutant of the same virus (148_STOP__116^myc^). Lysates collected at 4 dpi were subjected to immunoprecipitation (IP) using anti-myc antibody. (A) Cell lysates and (B) IP eluates were monitored by Western blot for the indicated proteins. n.s.: non-specific band(s).

## DISCUSSION

In this study, we have built upon our previous work that identified the an HCMV virion gH complex that lacks gL, and which instead contains UL116. Importantly, unlike gL, UL116 does not form a disulfide link to gH. Despite that our previous data indicate UL116 and gL compete for assembly onto gH, our findings here show that UL116 is required for high level expression of gH/gL complexes during infection. Therefore, we have demonstrated that UL116 plays a chaperone-like role for gH in HCMV. As illustrated in our model, we envision that UL116 stabilizes gH while gL forms a disulfide linkage to either UL128 or gO, and that UL148 further stabilizes gO, which we have previously shown is uniquely susceptible to ER-associated degradation (ERAD) (**FIG 11**)(24).

We further show that the gH/UL116 complex is abundant in HCMV virions (**FIG 7**). In fact, our results argue that gH/UL116 complex accounts for the long-noted observance of gH monomers in Western blot analyses of virion lysates resolved under non-reducing SDS-PAGE conditions. In such gels, gH present in gH/gL/gO (Trimer) and gH/gL/UL128-131 (Pentamer) complexes migrates as much larger species due to covalent disulfide linkages between gH and gL, and between gL and either gO or UL128 (20, 25). However, because UL116 does not form a disulfide link to gH, any gH/UL116 complexes in virions disassemble upon treatment with high concentrations of ionic detergents, such as SDS, causing the gH present in such complexes to migrate as an ~86 kD monomeric polypeptide species in SDS-PAGE.

Given its abundance in virions, whether gH/UL116 might play a role in viral entry remains an important unresolved question. Our data suggest that *UL116*-null viruses are poorly infectious, but the decreased expression of gH/gL complexes in such mutants is likely sufficient to explain this defect. Our finding of replication defects for *UL116*-null mutants contradict results from earlier genome profiling studies, which failed to note any substantial defects in viruses disrupted for *UL116* (11, 12). However, given that these profiling studies evaluated hundreds of mutants at once, and did not distinguish cell-free infectivity from cell-to-cell spread, the roughly ~10-fold replication defect we observed in the yield of cell-free infectious virus for such viruses disrupted in *UL116* may have gone unnoticed. Whether any replication defects of UL116 mutants reflect an inability to bind to hitherto yet-to-be identified cell surface to promote entry requires further investigation.

We also found evidence for an interaction between UL116 and UL148, which both appear to have a strong potential to affect the maturation of gH complexes and their abundance in virions. This suggests that regulation of cell tropism is likely far more complex than previously appreciated. We now know that UL148 activates the integrated stress response to remodel the endoplasmic reticulum, which is the site at which newly synthesized glycoproteins begin their journey to the cell surface or, for viral envelope glycoproteins, to TGN-derived vesicles in the juxtanuclear cytoplasmic assembly compartment where HCMV virions obtain their infectious envelopes. Although we previously reported that UL148 co-immunoprecipitates with gH, gL, UL131, and UL130 (6), subsequent studies in our laboratory indicate that of these, only the interaction with gH is repeatable (Siddiquey, M.N.A., Nguyen, C.C., and Kamil J.P., manuscript in preparation). Here, we observed that UL116 immunoprecipitates from infected cells contain UL148 and not gL, and these interactions could be validated in immunoprecipitation studies from transfected cells.

Together, our findings suggest interactions between, UL148, gH, and UL116 play roles in stabilizing immature forms of gH from being degraded (Fig. 10), which in turn may have effects on levels of gL, gO, and the Pentamer specific components UL128, UL130, and UL131. This model is supported by earlier work from our lab showing that gO is a constitutive ERAD substrate whose decay is attenuated by UL148 (24), as well as by our finding here that kifunensine, a chemical inhibitor of ERAD, causes a substantial increase in the steady state levels of gL but not gB during *UL116*-null but not wild-type virus infections.

**FIG 10.**
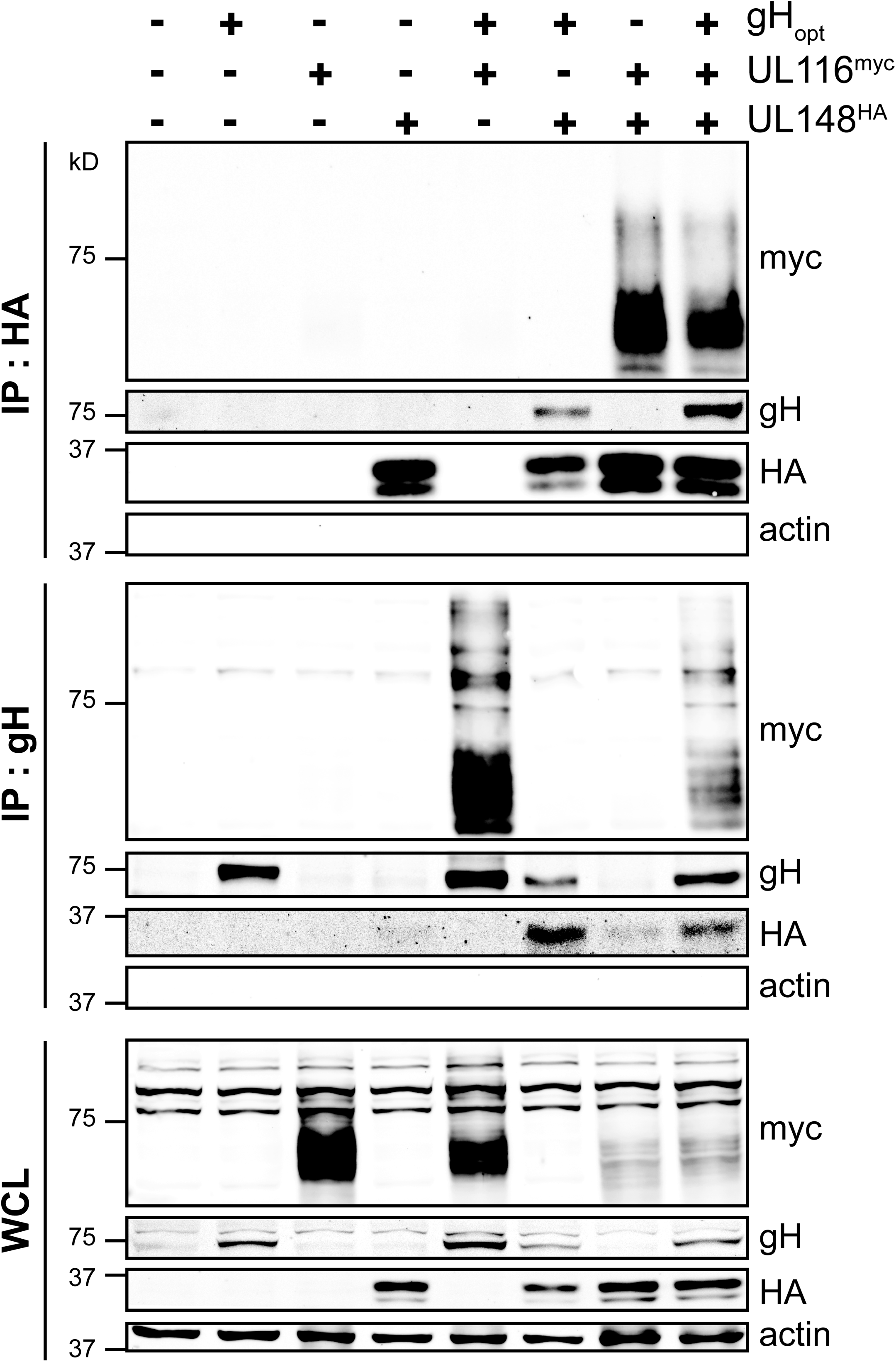
Reciprocal co-immunoprecipitation of gH, UL116, and UL148 from cells ectopically expressing the glycoproteins. HEK-293T cells in six-well cluster plates were transfected with 1 μg each of plasmid expression vectors encoding gH, UL116^myc^ or UL148^HA^; where appropriate empty vector was added such that equivalent amounts (3 μg) of plasmid DNA was transfected across all conditions. 48 h post transfection, cells were collected in lysis buffer and subjected to immunoprecipitation (IP) using anti-HA and anti-gH antibody. Cell lysates and IP eluates were monitored by Western blot for the indicated proteins.

**FIG 11.**
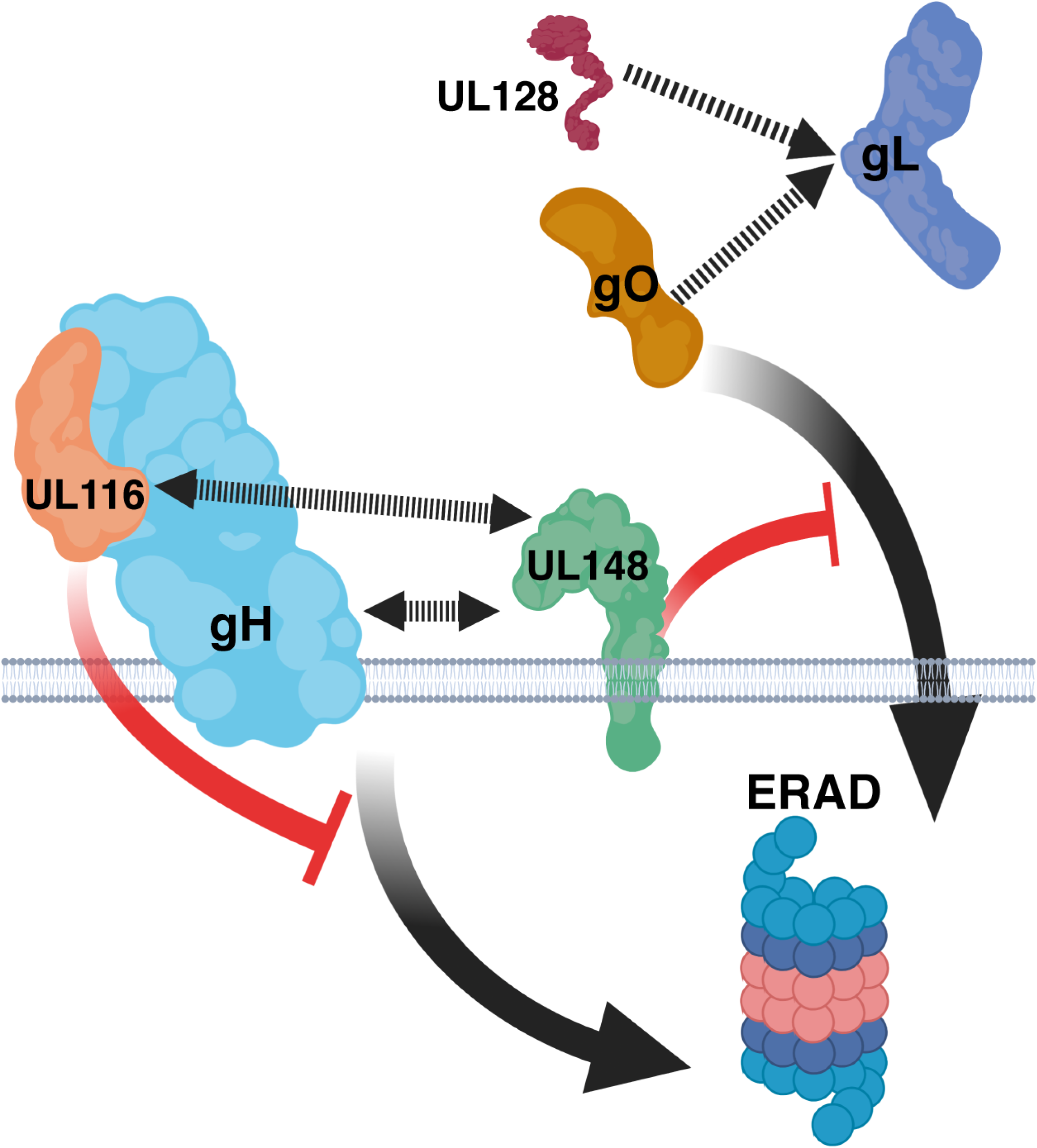
Model. UL116 acts as a gH chaperone, stabilizing gH while gL forms a disulfide bond to either gO or UL128. In the absence of UL116, increased amounts of gH are subjected to ER-associated degradation (ERAD). UL148, a viral ER resident glycoprotein, has been found to bind to gH and stabilizes gO from being targeted to ERAD. Our evidence of a physical interaction between UL116 and UL148 may suggest a functional relationship between the proteins.

UL116 is found to assemble onto gH at a site similar to that occupied by gL, which is consistent with it playing a role as a chaperone for gH, especially since UL116 does not covalently link to gH. gL, on the other hand, forms covalent links both to gH and to either UL128 or to gO. Hence, there may be good reason for UL116 to act as a placeholder to stabilize gH while gL forms disulfide linkages to either gO or UL128. Future work will no doubt be needed to test this model, and to decipher precisely how UL148 stabilizes gO (6, 24) and how its reported interactions with UL116 (this study), gH (6), the ERAD adaptor SEL1L (24), and the immune co-stimulatory ligand CD58 (26) may contribute to maturation of gH complexes. Regardless, because *UL116*-null mutants of strain AD169, which lacks UL148, are defective for expression of all gH complexes, the role of UL116 in stabilizing gH complexes does not appear to require UL148. Since UL116 appears to be more broadly conserved among betaherpesviruses than UL148 is, it will be no doubt interesting to determine whether evidence for gH chaperone functions can be found in other cytomegalovirus species.

## MATERIALS AND METHODS

### Cells

Telomerase-immortalized human foreskin fibroblasts (HFFT) derived from primary HFFs have been described elsewhere (24). ARPE-19 retinal pigment epithelial cells were purchased from ATCC (#CRL-2302), and human embryonic kidney 293T cells were purchased from Genhunter Corp. (Nashville, TN). Cells were cultured in Dulbecco’s Modified Eagle’s Medium (DMEM, Corning) supplemented with 25 μg/mL gentamicin (Invitrogen), 10 μg/mL ciprofloxacin (Genhunter), and either 5% fetal bovine serum (FBS, Sigma #F2442) or 5% newborn calf serum (NCS, Sigma #N4637). For experiments with Ad vectors done in the Ryckman lab (FIG 5), ARPE-19 were grown in DMEM/F12 medium (Gibco), supplemented with 10% FBS and penicillin-streptomycin, gentamicin, and amphotericin B.

### Viruses

Viruses were reconstituted from bacterial artificial chromosome (BAC) cloned virus genomes by electroporation of purified BAC DNA into HFFT. Wildtype HCMV strain TB40/E and its derivatives were reconstituted from TB40- BAC4 (27) or mutants thereof, and were grown on HFFT until 100% CPE was observed. For HCMV recombinants derived from AD169rv *rUL131*, a BAC clone of HCMV strain AD169 (28, 29) repaired for *UL131* (6, 24), viruses were amplified at low MOI in ARPE-19 cells until 100% cytopathic effect (CPE) was observed. Virus-containing culture supernatants were then subjected to centrifugation (1000 *× g)* for 10 min to pellet cellular debris. Cell-associated virus was then released by Dounce-homogenization of pelleted infected cells, clarified of cell debris by centrifugation (1000 ×*g*, 10 min), and combined with the cell-free medium then ultra-centrifuged (85,000 ×*g*, 1 h, 4°C) through a 7.5 mL 20% D-sorbitol cushion (25 mM Tris HCl pH 8.0, 1 mM MgCl_2_, 100 μg/mL bacitracin); resulting virus pellets were then resuspended in DMEM containing 20% NCS, or for analysis of virion glycoproteins, in Dulbecco’s phosphate-buffered saline lacking magnesium or calcium (DPBS, Lonza America, Inc.; Alpharetta, GA).

### New recombinant HCMV BACs and plasmids for this study

New recombinant viruses were constructed by *en passant* BAC recombineering (30, 31), as previously described (6, 18, 32–34). Briefly, for each recombinant virus, a primer pair (see below and Table 1) is used to PCR amplify an *I-SceI-AphAI* cassette from a BAC DNA template that contains the cassette. The cassette confers kanamycin resistance and is abutted by an I-Sce-I recognition site. The primers are designed to target homologous recombination into the targeted region of the BAC via bacteriophage lambda RecE/T and Gam (Red) recombinase activity, and also to generate ~40 bp repeats on each side of the inserted cassette, which allow it to later be excised during a second recombination step. The PCR product is electroporated into GS1783 *E. coli* (a gift of Gregory Smith, Northwestern University, Chicago, IL) carrying the BAC of interest to be modified. Kanamycin-resistant “integrate” colonies are isolated, and then subsequently resolved in a second step during which L-arabinose treatment is used to induce I-Sce-I homing endonuclease and a 41 °C heat shock is used to induce lambda Red recombinase activity. This step causes removal of the I-Sce-I *AphAI* kanamycin resistance cassette and leaves behind only the mutation, insertion, or deletion of interest. All primers and synthetic DNAs for this study were synthesized by Integrated DNA Technologies (Coralville, IA) and are shown in Table 1.

**TABLE 1.**
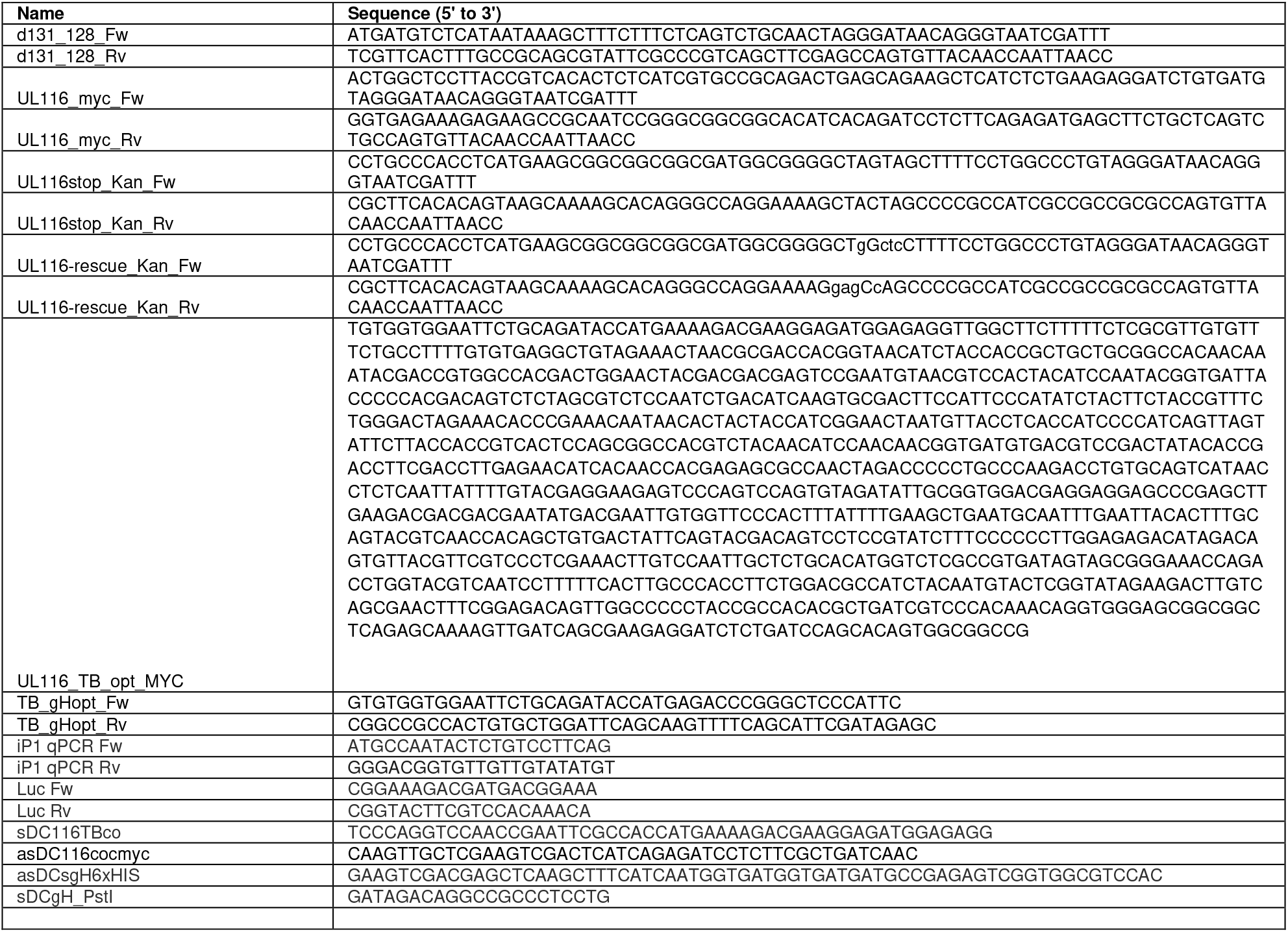
Primers and synthetic DNAs

To construct TB_116^myc^, a TB40-BAC4 derivative in which a Myc epitope tag was inserted at the C-terminus of *UL116*, the primer pair UL116_myc_Fw and UL116_myc_Rv was used to carry out en passant on TB40-BAC4. To construct TB_116_STOP_, in which tandem stop codons were inserted at *UL116*amino acid positions 10-11 in the context of TB40-BAC4, the primer pair UL116stop_Kan_Fw and UL116stop_Kan_Rv was used. A “rescue” virus was constructed from TB_116_STOP_ using the primer pair: UL116-rescue_Kan_Fw and UL116-rescue_Kan_Rv. TB_148_STOP__116^myc^, a strain TB40/E derivative null for *UL148* and also encoding a myc tag at the C-terminus of UL116, was constructed from TB_148_STOP_ (24), using UL116stop_Kan_Fw and UL116stop_Kan_Rv primers (Table 1). Similarly, an *AD169_rUL131* derived virus, AD169_rUL131_UL116_STOP_, was constructed using the primer pair UL116stop_Kan_Fw and UL116stop_Kan_Rv. For all new viruses, Sanger sequencing of modified regions was carried out by Genewiz, Inc. (Piscataway, NJ) to confirm the presence of intended changes without spurious mutations.

The plasmid pEF1_ UL116_GGS_Myc was constructed by inserting a synthetic dsDNA gBlock encoding a codon optimized (Homo sapiens codon bias) *UL116* ORF from strain TB40/E, fused to a Gly-Ser-Gly linker and a Myc epitope tag into the EcoRV site of pEF1 V5 His C (Invitrogen) using NEB HiFi Assembly Mix (New England Biolabs, Ipswitch, MA). All primers and synthetic DNAs for this study are shown in Table 1.

### Replication defective adenovirus (Ad) vectors

The nonreplicating (E1-) Ad vector expressing a soluble gH protein, in which the transmembrane and cytoplasmic UL75 (gH) domains were substituted by a hexahistidine (6xHis) tag sequence, was derived using a previously characterized AdMax shuttle vector expressing the full-length UL75 gH protein from HCMV strain TR (21). Briefly, a DNA fragment encompassing the downstream half of the codon optimized UL75 sequence was PCR amplified from the AdMax TR gH pDC316(io) shuttle plasmid (21) using primers sDCgH_PstI and asDCsgH6xHIS, resulting in an amplicon in which the 25 C-terminal amino acids of gH were replaced by a 6xHis tag. The amplicon was then ligated into the AdMax TR gH pDC316(io) using PstI-HindIII restriction sites. AdUL116myc was generated by subcloning the UL116_GGS_Myc sequence from pEF1_UL116_GGS_Myc plasmid to the AdMax pDC316(io) shuttle plasmid using the primers sDC116TBco and asDC116cocmyc (Table 1). AdUL115gL and AdGFP were described previously (21, 35). All Ad vectors were propagated and titered by TCID50 on HEK293IQ cells.

Human ARPE-19 epithelial cells or fibroblast cells (as indicated) were infected with each Ad vectors at 3 TCID50/cell, with the total Ad multiplicity standardized by the addition of Ad eGFP. Ad infected cultures were maintained in DMEM-F12 or DMEM with 2%FBS for 7 days. Supernatants were then harvested, adjusted to pH 8.0 using 1M Tris-HCl pH 8.0 and incubated at room temp for 1 h with Ni-NTA Agarose beads (Qiagen). Beads were washed twice with PBS, and bound complexes were eluted at room temperature for 20 min using 20 mM Tris-buffered saline (TBS) (pH 6.8) with 2% SDS, 10% glycerol, and bromophenol blue. Samples were resolved by nonreducing 6% SDS-PAGE, transferred to PVDF membranes in 25 mM Tris, 192 mM Glycine, and 10% methanol, pH 8.3. and probed with anti-gH (AP86, mouse monoclonal) or anti-gL (rabbit polyclonal) antibodies and detected using anti-rabbit or anti-mouse secondary antibodies conjugated to horseradish peroxidase (Sigma-Aldrich) and Pierce ECL-Western blotting substrate (Thermo Fisher Scientific). Chemiluminescence was detected using a Bio-Rad ChemiDoc MP imaging system.

### Virus titration

Infectivity of virus stocks and samples were determined by the tissue culture infectious dose 50% (TCID_50_) assay on hTERT immortalized fibroblasts, as previously described (18, 24, 36). Briefly, serial dilutions of virus were used to infect multiple wells of a 96-well plate. At 12 dpi, wells were scored as positive or negative for CPE, and TCID_50_ values were calculated according to the Spearman-Kärber method, as described previously (18).

### Virus growth kinetics

For determination of virus growth kinetics, fibroblasts (HFFT) and epithelial cells (ARPE-19) were seeded in a 24-well plate at 1.5 × 10^5^ cells per well. Cells were then infected at the indicated MOI in 0.5 mL of medium per well. Inocula were removed after 20 h, and the cells were washed two times with 1 mL of DPBS, and 0.5 mL of fresh DMEM 5% NCS was added to each well. Cell-free supernatants were collected at the indicated times post-infection and stored at −80°C until analysis. For determination of cell-associated virus titers at 5 dpi, cells were washed twice using 1 mL DPBS per wash, collected by scraping into 0.5 mL DMEM 5% NCS, and then stored at −80°C until analysis. After thawing, infected cell material was subjected to Dounce-homogenization and subsequently centrifuged (1000 ×*g*, 10 min) to pellet cell debris.

### Kifunensine treatment

Where indicated, kifunensine (KIF) (APeXBio) was applied at a final concentration of 2.5 μM at 48 hpi (MOI 1) for 24 h prior to collecting infected cell lysates at 72 hpi. Since the KIF stock solution was prepared at 2.5 mM in water, water (vehicle alone) was added to 0.1% in control treatments.

### Immunoprecipitation

For experiments with infected cells, a total of 1 × 10^7^ fibroblasts (HFFs) were infected at an MOI of 1 TCID_50_ per cell with TB_116^myc^ and TB_148_STOP__116^myc^ viruses. At 96 hpi, cells were washed once in DPBS and then lysed in lysis buffer (50 mM HEPES [pH 7.5], 1% Triton X-100, 400 mM NaCl, 0.5% sodium deoxycholate, 10% glycerol, supplemented with 1×Protease Inhibitor Cocktail (APeXBio Technology, LLC, Houston, TX). 250 μL of lysate were incubated with 2 μL of mouse Myc-Tag antibody clone 9B11 (Cell Signaling Technology, Cat. 2276S) for 4 h with rotation at 4°C. Twenty-five microliters of a 50% slurry of Protein G magnetic beads (Millipore Sigma, Cat. LSKMAGG10), pre-equilibrated in lysis buffer, were then added to each immunoprecipitation (IP) reaction, and reactions were allowed to rotate overnight at 4°C. The next day, the Protein G magnetic beads were washed three times in lysis buffer, and eluted by heating (50°C, 10 min) in 2×Laemmli buffer (4% SDS, 20% glycerol, 0.004% bromphenol blue, 0.125 M Tris HCl, pH 6.8) lacking reducing agent.

To obtain starting material for IP experiments from transfected cells, 3 μg of mammalian expression vector plasmids based on pEF1α V5 His C (Invitrogen) encoding codon-optimized *gH, UL116^myc^*, or *UL148^HA^* were transfected using 9 μL of TransIT-2020 reagent (Mirus, Inc, #MIR 5404) into 6 × 10^5^ HEK-293T cells that had been seeded into wells of a six-well cluster plate. As needed, transfections were supplemented with empty pEF1α V5 His C vector to maintain equivalent amounts of DNA per transfection reaction. Plasmid transfection reagent complexes were prepared and applied to cells according to the manufacturer’s instructions. At 48 h post transfection, cells were washed in DPBS and subsequently lysed in 200 μL of lysis buffer [50 mM HEPES (pH 7.5), 400 mM NaCl, 0.5%sodium deoxycholate containing 1×Protease Inhibitor Cocktail (APeXBio Technology, LLC, Houston, TX]. Ninety microliters of cell lysate were rotated at 4°C together with 10 μL of mouse anti-gH clone 14-4b antibody (37) for 4 h. Next, 25 μL of pre-equilibrated Protein G magnetic beads (as 50% slurry in lysis buffer) were added to each reaction, and rotated overnight at 4°C. Magnetic beads were then washed three times in lysis buffer and bound proteins were eluted as described above for infected cell IPs. For anti-HA IPs, 25 μL of anti-HA magnetic bead slurry (Pierce #8837), equilibrated in lysis buffer, were directly added to 90 μL lysate and incubated overnight before washing and processing for elution as described above.

### Tropism Studies

Fibroblasts (HFFTs) were infected at a MOI of 1 TCID_50_ per cell with TB_WT, TB_148_STOP_, and TB_116_STOP_ viruses. Duplicate aliquots of cell supernatants containing cell-virus were collected at 144 hpi. One aliquot was stored at −80°C for later infectivity analyses and the other aliquot was treated with DNAse I (RQ1 DNAse, Promega) prior to purification of viral DNA using a PureLink Viral RNA/DNA minikit (ThermoFisher), as described elsewhere (38). Viral genome copies eluted from the PureLink kit were then quantified using SYBR-green based real-time quantitative PCR (qPCR) assay using a pair of qPCR primers directed at intron A of the viral major immediate early locus: iP1 qPCR Fw and iP1 qPCR Rv (Table 1). The PCR efficiency of this primer pair was determined to be 99.4%. Fibroblasts (HFFT) and epithelial cells (ARPE-19) in 96-well plate format were then infected on the basis of viral genomes, using a 10-fold serial dilution series of virus genome equivalents. Each dilution of virus, starting with 1 ×10^6^ viral genomes per well and ending at 0.1 genomes per well, was applied to a row of eight wells of a 96-well cluster plate, with wells having been seeded at a density of 2 ×10^4^ cells per well. At 30 hpi, cells were fixed in ice-cold methanol, and stained using IE1 monoclonal antibody (mAb) clone 1B12, as described previously (39). To identify wells positive for virus, the aminoethyl carbazole (AEC) substrate kit (Thermo Fisher) was used to detect IE1 positive nuclei. Briefly, in this assay AEC staining detects IE1 mAb stained nuclei (a hallmark of HCMV-infected cells) by virtue of peroxidase activity from horseradish peroxidase-conjugated streptavidin bound to biotinylated secondary antibodies, which in turn are used to detect mouse anti-IE1 mAb 1B12. Infectivity was determined in units of TCID_50_ per viral genome in three biological replicates per experiment, and data normalized across different conditions by setting TB_WT infectivity on fibroblasts to 1.0.

### Western blots

Mouse mAb specific for UL116 (clone H4) was provided by GSK Vaccines (Rockville, MD). Mouse anti-gH mAb clone AP86 (40) was a generous gift from William J. Britt (University of Alabama, Birmingham). A custom generated anti-gL polyclonal antisera was raised in rabbits by immunization with synthetic peptide matching the C-terminal 21 amino acids of gL (Cys-KQTRVNLPAHSRYGPQAVDAR, gL residues 258-278) linked to keyhole limpet antigen via the appended cysteine residue at the N-terminus (Pacific Immunology, Ramona, CA). The peptide sequence used was identical to that used in a previous study to generate gL antisera (15). Western blot analyses of virion glycoproteins were carried out as described previously (6).

## ACKNOWLEDGEMENTS

This project was supported by NIH grants R01-AI116851 (to J.P.K), P30- GM110703 (LSUHS Center for Molecular and Tumor Virology), R01- AI097274 (B.J.R.), as well as an American Heart Association grant to E.P.S. (17POST33350043). Its contents are solely the responsibility of the authors and do not necessarily represent the official views of the funding agencies. We thank William J. Britt, Thomas E. Shenk, Klaus Osterrieder, B. Karsten Tischer, Gregory A. Smith, Christian Sinzger, and Ulrich Koszinowski for generously sharing reagents.

## Author Contributions

Performed research: M.N.A.S. except for FIG 5, which was done by Q.Y. Designed research: M.N.A.S., M.M., and J.P.K., except for FIG 5 was designed by E.J.S. and B.J.R. Interpreted data: M.N.A.S, E.P.S., B.J.R., M.M., J.P.K. Contributed new reagents, as follows: UL116 monoclonal antibody: D.A., G.V., D.Y., D.M., M.M. Contributed new adenovirus vectors: J.-M.L and B.J.R. Developed all other new reagents: M.N.A.S. and J.P.K. Obtained funding: J.P.K. and B.J.R. Wrote the manuscript: J.P.K., and B.J.R., with input from all authors.

